# DiI-mediated analysis of pre- and postsynaptic structures in human postmortem brain tissue

**DOI:** 10.1101/558817

**Authors:** Sujan C. Das, Danli Chen, W. Brandon Callor, Eric Christensen, Hilary Coon, Megan E. Williams

## Abstract

Most cognitive and psychiatric disorders are thought to be disorders of the synapse, yet the precise synapse defects remain unknown. Because synapses are highly specialized anatomical structures, defects in synapse formation and function can often be observed as changes in micro-scale neuroanatomy. Unfortunately, few methods are available for accurate analysis of synaptic structures in human postmortem tissues. Here, we present a methodological pipeline for assessing pre- and postsynaptic structures in human postmortem tissue that is accurate, rapid, and relatively inexpensive. Our method uses small tissue blocks from postmortem human brains, immersion fixation, DiI labeling, and confocal microscopy. As proof of principle, we analyzed pre- and postsynaptic structures from hippocampi of 13 individuals aged 4 months to 71 years. Our results indicate that postsynaptic CA1 dendritic spine shape and density do not change in adults, while presynaptic DG mossy fiber boutons undergo significant structural rearrangements with normal aging. This suggests that mossy fiber synapses, which play a major role in learning and memory, may remain dynamic throughout life. Importantly, we find that human CA1 spine densities observed using this method on tissue that is up to 28 hours postmortem is comparable to prior studies using tissue with much shorter postmortem intervals. Thus, the ease of our protocol and suitability on tissue with longer postmortem intervals should facilitate higher-powered studies of human pre and postsynaptic structures in healthy and diseased states.

## 1 Introduction

Neuropsychiatric disorders constitute a major health burden globally. Although the exact molecular mechanisms are unknown, altered neuronal connectivity is widely thought to underlie most psychiatric disorders (Penzes, Cahill, Jones, VanLeeuwen, & Woolfrey, 2011; Radley et al., 2008). Studies using advanced genetic methods including genome-wide association studies, whole genome sequencing, exome sequencing, and enrichment analysis indicate that genes altered in neuropsychiatric disorders frequently encode molecules that mediate synapse formation and function (Forrest, Parnell, & Penzes, 2018). Because synapse structure and function are tightly linked, alterations in synapse function often result in structural changes that can be observed using light microscopy. Some synaptic structures that can be affected include axonal and dendritic arborizations, presynaptic boutons, and postsynaptic dendritic spines. In particular, dendritic spines protruding from the dendritic shaft house critical postsynaptic elements of glutamatergic synapses. Electron microscopy indicates that the vast majority (>95%) of dendritic spines contain an asymmetric synapse (Arellano, Espinosa, Fairen, Yuste, & DeFelipe, 2007; Harris, Jensen, & Tsao, 1992). Moreover, spine size positively correlates with synapse size and strength (Berry & Nedivi, 2017; Harris, Fiala, & Ostroff, 2003). Because spines are large enough to be resolved by light microscopy, analysis of spine density and shape is a commonly used and reliable proxy for excitatory synapse density and function (Rochefort & Konnerth, 2012).

Although animal models are useful for studying psychiatric disorders, they cannot mimic human neuropathology to the full extent. Thus, human postmortem tissues provide an opportunity to study neuron and synapse morphology in psychiatric disorders. However, cellular and subcellular imaging in human postmortem tissues is hindered by the inability to rapidly fix human tissue by aldehyde perfusion. The most widely used method for analyzing synaptic structures in postmortem human brain tissue is Golgi staining but Golgi staining typically uses two dimensional imaging and can underestimate spine density by as much as three-fold (Shen, Sesack, Toda, & Kalivas, 2008). Electron microscopy is the gold standard for direct synapse analysis but it is expensive, time-consuming, and requires specialized technical skills and reagents. Single cell microinjection of fluorescent dyes followed by confocal microscopy generates spectacular high-resolution, three-dimensional images of single neurons (Benavides-Piccione, Fernaud-Espinosa, Robles, Yuste, & DeFelipe, 2013; Merino-Serrais et al., 2013). However, even in mice, microinjection can be a fickle method that requires a precise level of fixation, does not label axons well, and works best in human tissues with very short postmortem intervals. Taken together, all of these methods of human brain analysis suffer from very low throughput, which generates underpowered studies. Thus, the field is in need of an easy, inexpensive, and higher throughput method of analyzing pre- and postsynaptic structures in the human brain.

Here, we demonstrate that neurons from small pieces of postmortem human brain tissue, which are more easily acquired from autopsy than whole brains, can be labeled and imaged at the resolution of dendritic spines and presynaptic boutons using the lipophilic dye, DiI. DiI labeling is fast, inexpensive, and does not require special equipment yet generates high-resolution three dimensional images of neuron morphology across a wide range of tissue ages and postmortem intervals. We demonstrate that DiI is suitable for labeling most neuronal structures including axons, dendrites, spines, and presynaptic boutons. As proof of concept, we tested for age-dependent changes in hippocampal connectivity using 13 control brains across the lifespan. Interestingly, we find dendritic spine density and shape in hippocampal CA1 neurons is remarkably stable with age. In contrast, DG mossy fiber presynaptic structure shows age-dependent structural plasticity. To our knowledge, this is the most extensive analysis of CA1 spine density and the first analysis of DG mossy fiber boutons in human brain tissue. In summary, our newly proposed DiI-based labeling of postmortem human brain tissue should enable higher-powered studies aimed at elucidating changes in neuron morphology associated with neuropsychiatric disorders.

## 2 Methods and Materials

### 2.1 Tissue preparation

Human hippocampal tissue blocks were obtained through Utah’s State Office of the Medical Examiner (OME). The brain tissue collection procedures are approved under an exemption (45CFR46.102(f)) from the Institutional Review Board (IRB) at the University of Utah. The age, sex, postmortem interval, and manner of death for each case is summarized in Table

1. We used postmortem hippocampus samples from 13 sudden death cases (10 males and 3 females). All the cases tested negative for acetone, isopropanol, methanol, cocaine, methamphetamine, morphine, and THC in toxicology reports. Upon brain removal for autopsy at the OME, a 1.5 cm^3^ hippocampal block was immediately immersed in cold 4% (w/v) paraformaldehyde in 0.1 M Phosphate buffer (PB) (pH 7.4). After 1 hour of fixation, the hippocampal block was post-fixed in cold 4% paraformaldehyde plus 0.125% (v/v) glutaraldehyde in PB (pH 7.4) for 24 hours at 4°C. Note: this fixation time is optimized for 1.5 cm^3^ postmortem human tissue blocks. Tissue blocks with different sizes may require different fixation times. After post-fixation, 350 µm coronal sections were cut on a vibratome (Leica VT1200). Fixed brain slices may be stored in PB at 4°C for at least one month for DiI based labeling of dendritic spines and mossy fiber boutons.

### 2.2 DiI-labeling

Microscopic 1,1’-dioctadecyl-3,3,3’,3’-tetramethylindocarbocyanine perchlorate (DiI) crystals (ThermoFisher D282) with a particle diameter of approximately 10-20 µm were placed in the CA1 sub-region and supra-pyramidal blade of the DG for dendritic spine and mossy fiber bouton analysis, respectively. A brain section was placed on a glass slide and the surface liquid was gently removed with a laboratory tissue. DiI crystals were manually placed just under the tissue surface using a pulled capillary glass (World Precision Instruments, 1B120F-4) using a dissecting microscope. Slices with the DiI crystals were then placed in PB (pH 7.4) in a 12-well plate and incubated at 37°C. After 72 hours, sections were checked under a fluorescent microscope for sparse labeling of CA1 neurons and DG axons. Tissue sections with DiI labeling were coverslipped in PB and confocal imaging was performed within 48 hours of coverslipping.

### 2.3 Confocal imaging, reconstruction, and analysis

Confocal imaging was performed on the CA1 and CA3 regions of the hippocampus for dendritic spines and mossy fiber boutons, respectively. Images were acquired with a Zeiss LSM 710 confocal microscope using a 63X oil immersion objective (NA=1.4). For both basal and apical dendrites, we imaged secondary dendrites of comparable caliber (mean basal dendrite area-to-length ratio: 2.62±0.04 (SEM); mean apical dendrite area-to-length ratio: 2.72±0.07 (SEM)) located 70-100 µm from the soma. All images were obtained with a cubic voxel of 0.07 µm^3^. Images of dendritic segments and mossy fiber boutons were deconvolved using AutoQuant X3 (Bitplane, RRID: SCR_002465) using the following settings: total iterations: 10, save interval: 10, noise level: low, and noise value: 2. Three dimensional spine and bouton modeling was performed using Neurolucida 360 (MBF Bioscience). The average length of analyzed dendritic segments ranged from 17.11 µm to 36.19 µm. Dendritic segments and spines were traced using Rayburst Crawl tracing method and manually inspected for errors. After modeling, spines were classified into mushroom, stubby and thin categories based on metrics previously described (Rodriguez, Ehlenberger, Dickstein, Hof, & Wearne, 2008). Spine parameters such as spine head diameter (HD), spine length (L) and neck diameter (N) were used to classify the spines. Spine classification settings are- mushroom [head-to-neck ratio (HNR) >1.1 and HD > 0.35µm], thin [head-to-neck ration (HNR)>1.1 and HD <0.35 µm; HNR<1.1 and length-to-head ratio >2.5], stubby [head-to-neck ratio (HNR) < 1.1 and length-to-head ratio <2.5]. Mossy fiber boutons were traced with soma function using a 12 µm interactive search region and 1 µm size constraint. Filopodia were then traced using the Rayburst Crawl method.

### 2.5 Statistics

Statistical analyses were performed in Prism (GraphPad Software) and the SAS software package (www.sas.com). To test whether age or postmortem interval affect spine and mossy fiber bouton parameters, we used mixed effect models with post hoc analyses. A mixed effects model estimates the independent effects of predictors (age and postmortem interval) accounting for clustering in the data, because multiple spine and bouton parameters were measured from the same neuron within cases, and additionally multiple neurons were measured for the same case. This clustering can lead to inflated significance, as measures may be more similar within the same neuron, and as neurons may be more similar within the same case. Our mixed effect model therefore treated age, and postmortem interval as fixed effects, and neurons (within cases) and the cases as random effects. Analyses were done within dendrite type (apical or basal), and differences between dendrite type was done using two-tailed unpaired t-tests. Finally, for any parameter where overall significance for age was achieved with the basic model, we explored this significance in two ways. First, we testing if a non-linear age model more accurately explained the data through including an additional age-squared term. Second, we tested for a binary effect of very young cases (two years old or younger) versus all other cases by creating a binary young-vs.other age variable, then investigating least squares means (means adjusted for other model predictors) of the parameters within these two age groups.

### 2.6 Data sharing statement

Raw data including images and 3D models will be made available upon request.

## 3 Results

### 3.1 A simple methodological pipeline for analyzing neuron morphology in human postmortem tissue

We sought to develop a relatively fast, low cost, and easily accessible method for high resolution three-dimensional neuron labeling and imaging in human postmortem tissue. We based our method on the fluorescent, lipophilic dye, DiI. DiI only fluoresces when it is incorporated into a cell membrane and it freely diffuses among a labeled cell membrane without crossing between adjacent or connected cells. These are ideal properties for labeling and imaging the intricate details of individual neurons. We collected 1.5 cm^3^ pieces of hippocampal brain tissue from de-identified subjects at the time of autopsy (Figure 1a). The tissue is immediately immersed in ice cold paraformaldehyde for 24 hours. The tissue block is then cut into 350 µm sections (Figure 1b) and microscopic DiI crystals (10-20 um in diameter) are placed in the region of interest using a glass micropipette and a dissecting microscope (Figure 1c,d). For dendrite labeling and short range axon labeling within the hippocampus, DiI diffusion is complete within 72 hours (Figure 1e). Sections are then mounted for confocal imaging. Confocal Z-stacks through the region of interest are collected, deconvolved, and modeled in three dimensions using Neurolucida 360 software (Figure 1f-h). It should be noted that one caveat of this method is that fluorescent DiI labeling is not permanent. Imaging should be completed within 5 days of placing the DiI crystal or else the labeling will become blurry in the paraformaldehyde fixed and, thereby, lightly permeabililized tissue. However, the fact that this method generates high quality neuron labeling from small blocks of immersion fixed brain tissue using no specialized reagents or technical skills should make this method feasible for most neuroscience laboratories with access to postmortem tissue.

**Figure 1:**
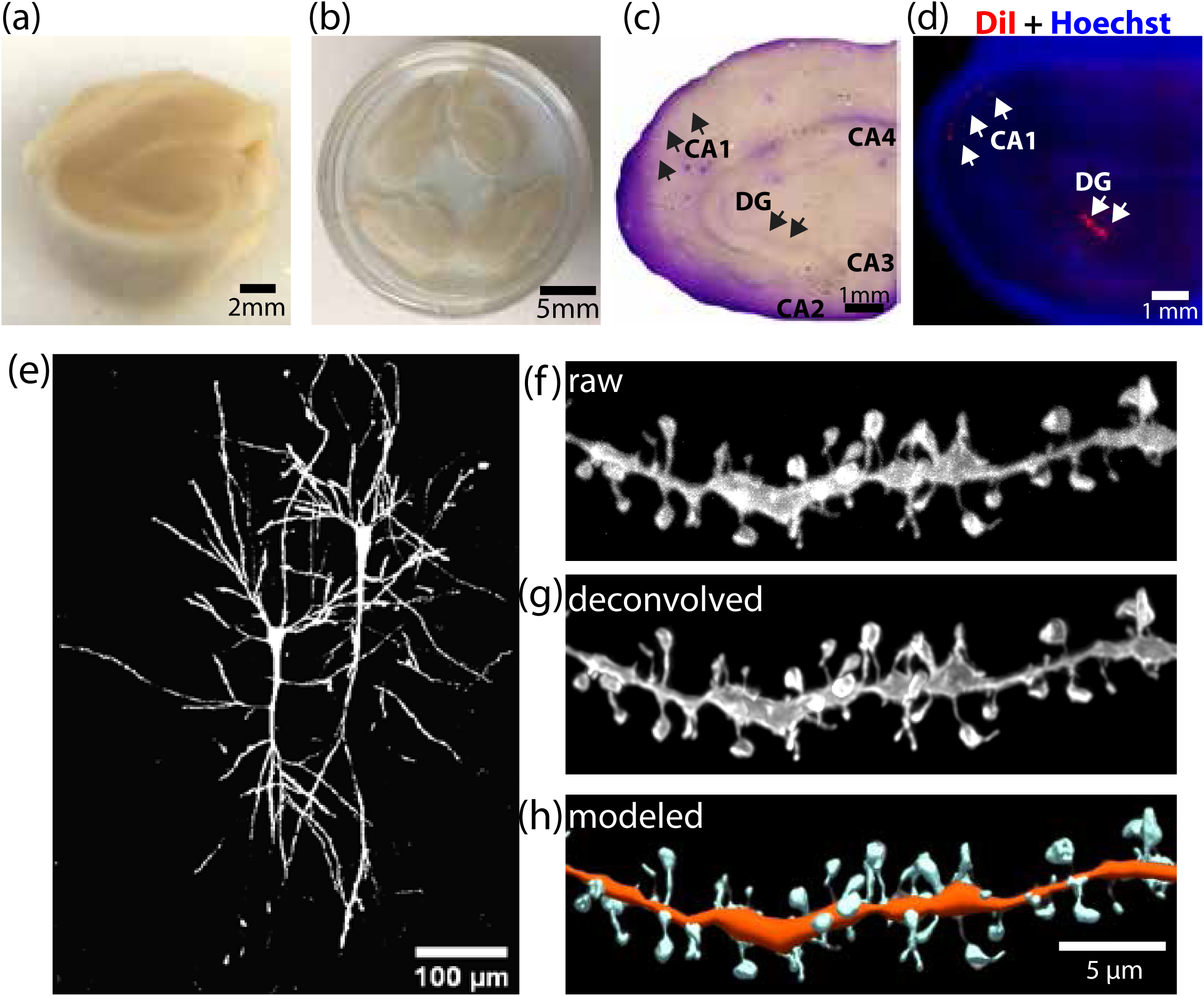
Methodological pipeline for DiI-labeling and spine imaging from postmortem human tissue. (a) Starting material: a ∼1.5 cm^3^ piece of human hippocampus. (b) 350 µm coronal slices of human hippocampus. (c) A Nissl-stained section of human hippocampus to reveal sub-regions as indicated. Arrows indicate region of placement for DiI microcrystals (d) Fluorescent image of section shown in (c). DiI crystals are red and cell nuclei are labeled with Hoechst. (e) Tiled maximum z projection of a confocal image of a CA1 neuron labeled with DiI. (f) Maximum z projection of a confocal image of a CA1 dendrite labeled with DiI. The raw confocal image is shown. (g) The dendrite in (f) was deconvolved. (h) The deconvolved image from (g) was used to generate a 3D model for analysis. CA, Cornu Ammonis; DG, Dentate Gyrus.

### 3.2 Analyzing synaptic structures underlying learning and memory

The hippocampus is a critical brain region necessary for many forms of learning and memory (Broadbent, Squire, & Clark, 2004). Thus, we examined whether morphological differences in hippocampal synaptic structures could be observed in tissue from healthy subjects of various ages across the lifespan. We started by analyzing spine density in CA1 pyramidal neurons. It is well established that spine number and morphology provide a proxy for excitatory synapse density (Hering & Sheng, 2001) and CA1 spines are a major site of long-term potentiation, which is widely regarded as the cellular mechanism underlying learning and memory (Whitlock, Heynen, Shuler, & Bear, 2006), yet CA1 spines have not yet been extensively analyzed in postmortem human tissue.

Human CA1 neurons from postmortem tissue were labeled with DiI. 2-4 segments of second order apical dendrites were imaged per neuron and 2-4 neurons were imaged for each brain. Figure 2 shows a representative set of neurons and dendritic segments from several brains. Note that despite being from tissue fixed 11-18 hours postmortem, we do not observe blebbing, neurite fragmentation, or other obvious hallmarks of cell death, hypoxia, or ischemia (Figure 2). Though we cannot exclude that any postmortem interval leads to changes in neuron morphology, the neuron morphology in our samples appears well preserved and the secondary dendrites are a consistent caliber with even DiI labeling and diffusion. Qualitatively, we find that dendrite and spine labeling is highly consistent from different dendrites within one neuron (Figure 2b-g) and from different brains (Figure 2h-m).

**Figure 2:**
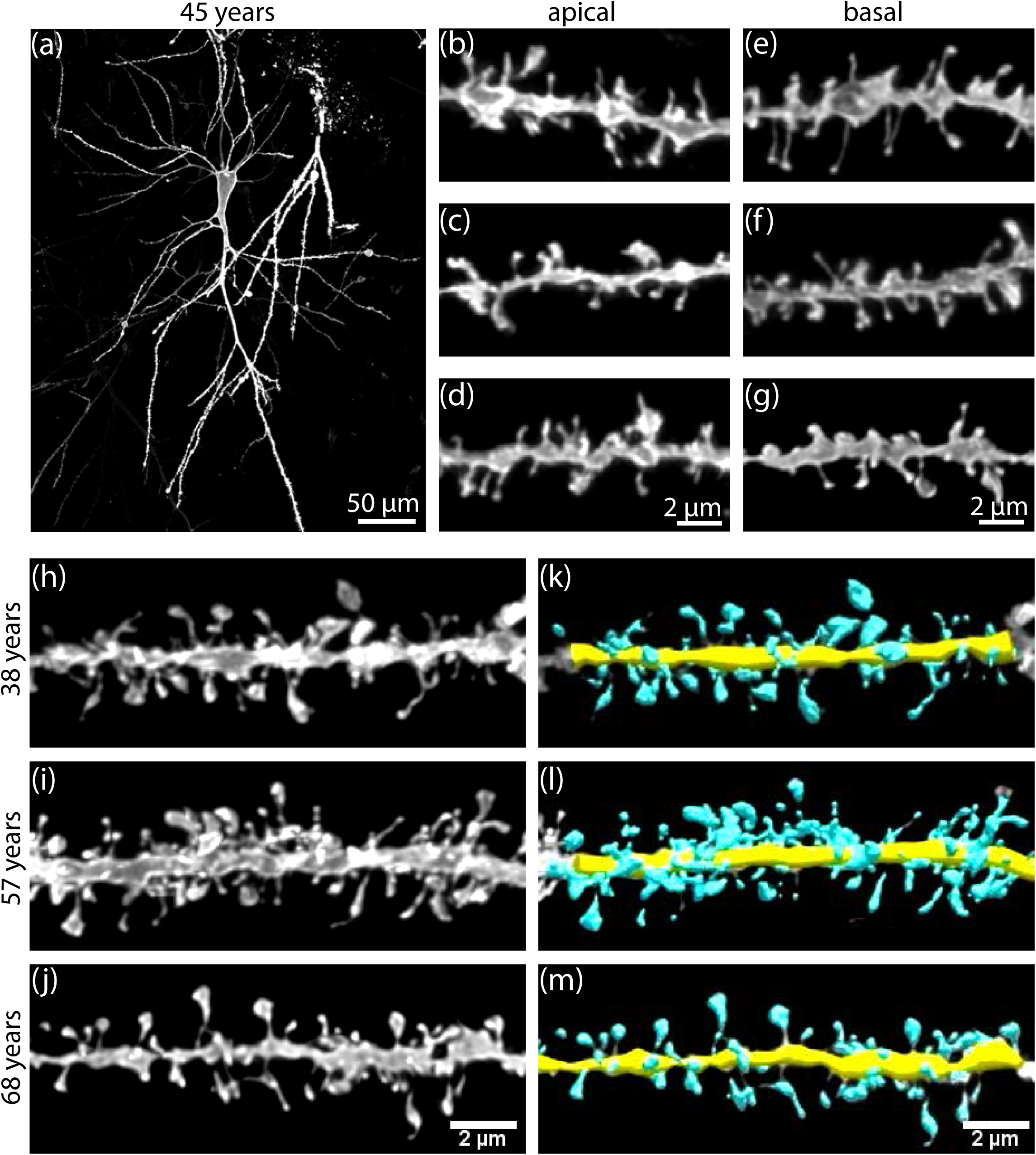
Consistent DiI labeling of dendritic segments across different neurons and brains. (a) 20x tiled magnification image of a DiI-labeled CA1 pyramidal neurons. (b-g) Examples of apical and basal dendritic segments used for analysis from the labeled neuron shown in (a). (h-j) Apical dendritic segments from 3 different brains of indicated ages that were used for analysis. (k-m) 3D Neurolucida models of the apical dendritic segments shown in (h-j). All images are maximum z projections and (b-j) show deconvolved maximum z projections of confocal images.

### 3.3 CA1 dendritic spine parameters are stable in adults across a range of ages and postmortem intervals

Next, we sought to expand our data set and test if CA1 spine parameters vary with age or postmortem interval. We analyzed spine density and morphology from 11 cases with varying age from 5 months to 71 years old and postmortem intervals from 10 to 28 hours (Table 1). We also expanded our analyses to include CA1 apical and basal dendrites because spine morphology and functional synapse properties have been shown to differ between CA1 apical and basal dendrites in rodents (Basu et al., 2017; Haley, Schaible, Pavlidis, Murdock, & Madison, 1996) but have not been compared in humans.

**Table 1.** Demographics of the cases.

| Sample ID | Age | Sex | Postmortem interval (Hours) | Manner of Death |
| --- | --- | --- | --- | --- |
| U76<sup>*</sup> | 4 Months | Male | 25.5 | Pending |
| U66 | 5 Months | Female | 15 | Accident |
| U51<sup>#</sup> | 2 Years | Female | 18 | Homicide |
| U78 | 23 Years | Male | 12 | Accident |
| U82 | 27 Years | Female | 10 | Accident |
| U71 | 38 Years | Male | 18 | Accident |
| U56 | 45 Years | Male | 11 | Natural |
| U58<sup>*</sup> | 57 Years | Male | 18 | Natural |
| U63 | 57 Years | Male | 18 | Natural |
| U41<sup>#</sup> | 58 Years | Male | 13 | Natural |
| U61<sup>#</sup> | 68 Years | Male | 18 | Accident |
| U74 | 70 Years | Male | 15.5 | Accident |
| U67 | 71 Years | Male | 28 | Natural |
| <sup>#</sup>cases used for CA1 spines analysis only; <sup>*</sup>cases used for mossy fiber bouton analysis only. |  |  |  |  |

Our results indicate that CA1 apical spine density, spine head width, and spine length is remarkably similar among all subjects analyzed (Figure 3a,d,g). The same is true for basal dendrites (Figure 3b,e,h). We used a mixed effect model to statistically test if apical or basal CA1 spine density or morphology varies with age or postmortem interval (Table 2). This analysis indicates that the postmortem intervals of our tissue from 10 to 28 hours do not significantly affect either apical or basal spine density (apical, p=0.2808; basal, p=0.9276), spine head diameter (apical, p=0.1818; basal, p=0.8164), or spine length (apical, p=0.3885; basal, p=0.8732) (Table 2). Similarly, we found that age does not affect any parameter except apical spine density (p=0.0324) (Table 2). Inspection of the distribution of apical spine density by age suggested slight differences for cases ≤ 2 years old. A post-hoc analysis excluding cases ≤ 2 years old indicates no significant effect of age (p=0.7117) on apical spine density. The least squares mean apical spine density for measurements from cases ≤ 2 years old was 3.21, which is significantly higher than the least squares mean of 2.24 found for the rest of the age distribution (p<0.0001). In summary, we find that CA1 apical spine density is greater in very young brains but that in adults, CA1 spine density is stable across ages.

**Table 2.** Mixed model analyses to determine effects of age and postmortem interval (PMI) on spine characteristics, adjusting for multiple neurons measured on each case.

| Spine Density |  |  |  |  |
| --- | --- | --- | --- | --- |
| Variable | Basal |  | Apical |  |
|  | Estimate | p-value | Estimate | p-value |
| Age | -0.0024 | 0.6476 | -0.0125 | 0.0324 |
| PMI | 0.0025 | 0.9276 | 0.0325 | 0.2808 |
| Spine Head Diameter |  |  |  |  |
| Variable | Basal |  | Apical |  |
|  | Estimate | p-value | Estimate | p-value |
| Age | 0.0002 | 0.6858 | -0.00004 | 0.8802 |
| PMI | -0.0006 | 0.8164 | 0.0019 | 0.1818 |
| Spine Length |  |  |  |  |
| Variable | Basal |  | Apical |  |
|  | Estimate | p-value | Estimate | p-value |
| Age | -0.0008 | 0.6234 | -0.0016 | 0.1562 |
| PMI | 0.0015 | 0.8732 | -0.0052 | 0.3885 |

**Figure 3:**
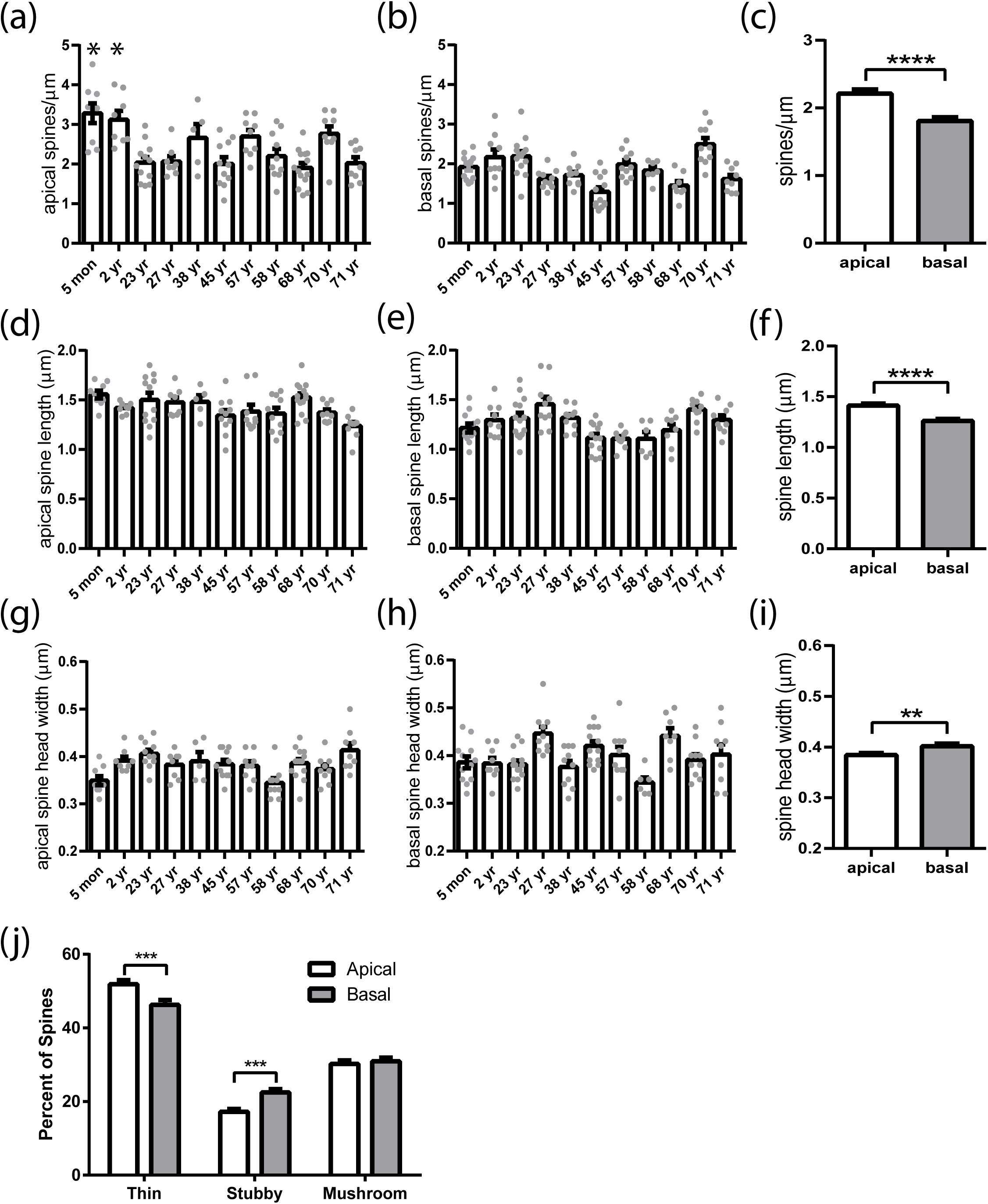
Human CA1 spine morphology differs between apical and basal spines but does not change with adult age. (a) Bar graph of mean CA1 apical spine density for each brain examined. Each point represents one dendritic segment. n=5-14 dendritic segments per brain. Statistics were analyzed using a mixed effects ANOVA model to account for nested data and are shown in Table * indicates that the spine density in the 5 month and 2 years old brain is slightly yet significantly higher than all other ages. (b) The same as in (a) but for CA1 basal spines. n=5-14 dendritic segments per brain. No differences are significant. (c) Bar graph comparing the mean CA1 apical and basal spine density in adult brains. n=87-91 dendritic segments per group, two-tailed unpaired t-test, p<0.0001. (d) Bar graph of mean CA1 apical spine length for each brain examined. Each point represents one dendritic segment. n=5-14 dendritic segments per brain. No differences are significant. (e) The same as in (d) but for CA1 basal spines. n=6-14 dendritic segments per brain. (f) Bar graph comparing the mean CA1 apical and basal spine length in adult brains. n=87-90 dendritic segments per group, two-tailed unpaired t-test, p<0.0001. (g) Bar graph of CA1 apical spine head width for each brain examined. Each point represents one dendritic segment. n=5-14 dendritic segments per brain. No differences are significant. (h) The same as in (g) but for CA1 basal spines. n=6-14 dendritic segments per brain. No differences are significant. (i) Bar graph comparing the CA1 apical and basal spine head width in adult brains. n=86-90 dendritic segments per group, two-tailed unpaired t-test, p=0.0086. (j) Bar graph comparing spine categories as a percentage of total spines between CA1 apical and basal dendritic segments. ordinary two-way ANOVA followed by Bonferroni’s multiple comparison test, n=87-91 dendritic segments per group. All data in all graphs represented as mean ± SEM. *, p<0.05; **, p<0.01; ****p<0.0001.

Rodent models suggest that spine morphology and the synaptic strength of CA1 basal and apical dendrites is mediated through different molecular pathways (Basu et al., 2017; Brzdak et al., 2019). Thus, we tested if structural differences are present between CA1 apical and basal spines in human hippocampus. Given that we did not find an effect of adult age or postmortem interval on either apical or basal spine parameters, we pooled adult cases to directly compare apical and basal spine parameters. Here, we found some significant differences between apical and basal spine morphology. Specifically, we find that apical spines have a significantly higher average spine density (in spines/µm ±s.e.m. apical=2.21±0.06, basal=1.81±0.05, p<0.0001) and spine length (in µm ±s.e.m. apical=1.41±0.02, basal=1.26±0.02, p=0.0001) but slightly smaller spine head diameter (in µm ±s.e.m. apical=0.38±0.003, basal=0.40±0.005, p=0.0086) than basal spines (Figure 3c,f,i). Next, we classified spines into thin, stubby and mushroom categories (see methods). These common spine morphology categories take into consideration spine length, head width, and neck width (see methods) and are known to provide information about the maturity of the synapse. In particular, mushroom shaped spines, which have a high spine head width to neck ratio, are considered the most mature or potentiated synapses (Hering & Sheng, 2001). We find that in adult humans, CA1 apical dendrites have slightly more thin spines (in % ±s.e.m. apical=51.89±1.061, basal=46.30±1.27, p=0.0003) and fewer stubby spines (in % ±s.e.m. apical=17.24±0.76, basal=22.46±0.91, p=0.0009) than basal dendrites but the percent of mushroom spines (in % ±s.e.m. apical=30.21±0.95, basal=30.91±1.04, p>0.99) is not significantly different (Figure 3j). Together, this analysis indicates that CA1 apical and basal spines tend to have slightly different spine parameters but when more complex measures of spine shape are considered, CA1 apical and basal dendrites likely have a similar ratio of mature synapses in adult humans.

### 3.4 Hippocampal mossy fiber bouton morphology changes with age in humans

The hippocampus also contains unmyelinated axons of dentate granule (DG) cells, called mossy fibers, which form unique, multi-neuronal synaptic structures with CA3 pyramidal neurons and nearby GABA neurons. These so called mossy fiber synapses consist of a large main bouton that synapses onto CA3 pyramidal neurons and synaptic projections, called filopodia, that originate from the main bouton to form synapses with GABAergic interneurons (Figure 4a). While the main synapse provides strong mono-synaptic excitation to CA3 neurons, filopodia synapses excite GABA neurons that, in turn, inhibit CA3 neurons (Scharfman, 2016). This feed-forward inhibition mediated by mossy fiber filopodia is necessary for normal learning and memory in mice (Guo et al., 2018; Ruediger et al., 2011) but mossy fiber filopodia have never been examined in humans.

**Figure 4:**
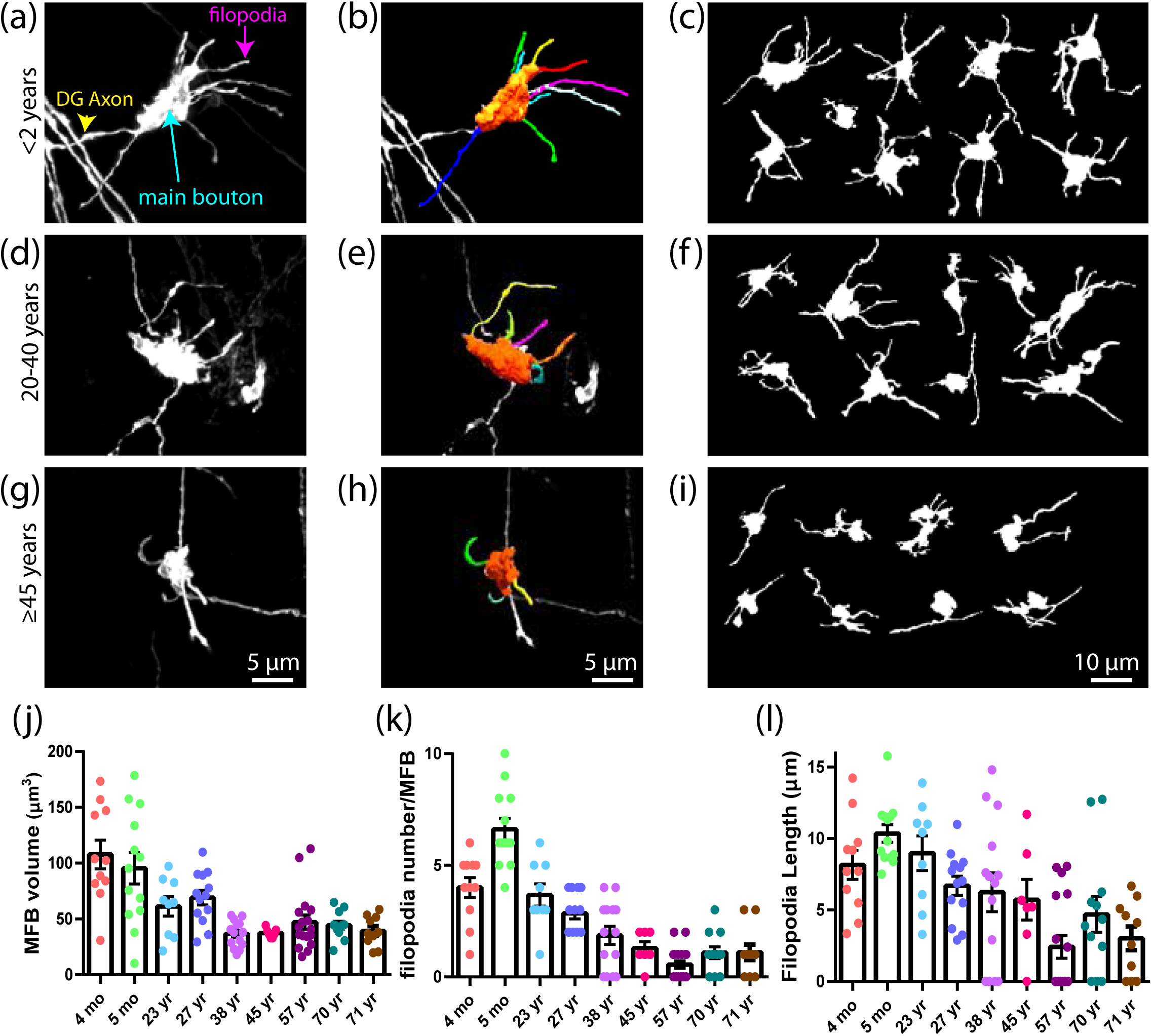
DG mossy fiber bouton morphology changes with age in human post-mortem tissue. (a, d, g) Deconvolved images of mossy fiber boutons from human postmortem tissue. (b, e, h) 3D Neurolucida 360 models of mossy fiber boutons shown in (a, d, g). (c, f, i) Tracings of maximum z projections from confocal images of mossy fiber boutons. Shown are numerous representative boutons from brains in the indicated age ranges of <2 years, 20-40 years and ≥45 years. (j) Bar graph of mean mossy fiber bouton volume per brain. n=5-14 boutons per case. (k) Bar graph of mean filopodia number per mossy fiber bouton per brain. n=5-14 boutons per case. (l) Bar graph of mean filopodia length per mossy fiber bouton per brain. n=5-14 boutons per case. In (j-l), each age is represented by one brain and each dot represents one mossy fiber bouton. All data represented as mean±SEM. Statistics were analyzed using a mixed effects model to account for nested data and are shown in Table 3. Age but not postmortem interval is a significant factor for each analysis.

Here, we tested if presynaptic mossy fiber structures can be labeled and analyzed with DiI in human postmortem tissues. We find that DiI clearly labels individual DG axons and their associated mossy fiber synapse complexes in sections of immersion fixed human brain tissue (Figure 4a,d,g) and they can be modeled in three dimensions using Neurolucida360 (Figure 4b,e,h) similar to dendritic spines. We imaged and analyzed the number of filopodia, filopodia length, and volume of the main bouton for 107 mossy fiber boutons from 10 brains (5-14 boutons/brain) (Table 1). We found that the volume of the mossy fiber bouton, density of filopodia per bouton, and filopodia length all decrease with age in humans (Figure 4). As we did for CA1 spines, we used a mixed model analysis to test if age or postmortem interval have a statistically significant impact on mossy fiber bouton structure (Table 3). Similar to CA1 spine morphology, we found no significant effect of postmortem interval on filopodia number (p=0.9678), filopodia length (p=0.3155), or bouton volume (p=0.2515) (Table 3). However, unlike CA1 spines, age has a significant effect on filopodia number (p=0.0013), filopodia length (p=0.0011), and bouton volume (p=0.0018) (Table 3). Interestingly, close inspection of the distribution of mossy fiber bouton and filopodia density by age show a slight increase in filopodia density in the oldest ages (Figure 4j,k) and suggest the effects of age may be non-linear. Additional analyses including a nonlinear age*age effect showed significance for filopodia number and bouton volume (Table 3). These results account for the increase in both of these parameters in measurements from both the oldest and youngest cases.

**Table 3.** Mixed model analyses of age and postmortem interval (PMI) effects on mossy fiber bouton parameters.

| Filopodia Number |  |  |
| --- | --- | --- |
| Variable | Estimate | p-value |
| Age | -0.0668 | 0.0013 |
| PMI | 0.0024 | 0.9678 |
| Filopodia Length |  |  |
| Variable | Estimate | p-value |
| Age | -0.0894 | 0.0011 |
| PMI | -0.0825 | 0.3155 |
| Mossy Fiber Bouton Volume |  |  |
| Variable | Estimate | p-value |
| Age | -0.8902 | 0.0018 |
| PMI | 1.0152 | 0.2515 |
| Additional models allowing for non-linear effects of age. |  |  |
| Filopodia Number |  |  |
| Variable | Estimate | p-value |
| Age | -0.1728 | 0.0049 |
| Age*Age | 0.0015 | 0.0341 |
| PMI | -0.0932 | 0.1359 |
| Mossy Fiber Bouton Volume |  |  |
| Variable | Estimate | p-value |
| Age | -2.2548 | 0.0087 |
| Age*Age | 0.0200 | 0.0536 |
| PMI | -0.2222 | 0.7930 |

## 4 Discussion

### 4.1 A DiI-based method for analyzing neuron structure in human postmortem tissues

Pre- and postsynaptic structures are the physical traces of synapses and provide clues to synaptic strength and connectivity yet analysis of synaptic structures in human brain tissue is limited due to cumbersome methodology required. Here, we describe a DiI-based labeling of dendritic spines and axon boutons in human postmortem tissue samples and use it to investigate pre- and postsynaptic structures in the healthy human hippocampus across the lifespan. This method has several advantages over existing methods. First, it is highly accurate yet faster, easier, and less expensive than common methods. Second, it can be done on human tissue samples within a range of postmortem intervals (tested up to 28 hours here), enabling larger sample sizes and higher-powered studies. Third, post-fixed tissue slices can be stored up to a month prior to placing DiI crystals, providing a convenient time window for labeling. Fourth, although we analyzed pre- and postsynaptic structures in the hippocampus, this method should be easily adapted to any other brain region. Overall, our proposed DiI-labelling should benefit researchers for the fast and accurate study of neuronal connectivity in healthy and disease conditions using human postmortem tissues.

### 4.2 DiI labels intact neurons in human postmortem tissue from a range of postmortem intervals

The requirement for extremely short postmortem intervals to obtain high quality three dimensional spine reconstructions severely limits tissue collection and results in studies with low sample sizes, usually a few neurons from 1-3 brains (Benavides-Piccione et al., 2013; Merino-Serrais et al., 2013). In contrast, our DiI-based methodological pipeline generates high quality dendritic spine labeling in human tissues with a longer postmortem delay; thus far we tested up to 28 hours postmortem. When labeled, the neurons do not show obvious morphological signs of hypoxia, ischemia, or damage cause neuron such as blebbing, fragmentation, or poor diffusion of DiI (Fig 1e), suggesting good membrane integrity and relatively healthy tissue. Here, we report a mean CA1 apical spine density of 2.21 spines/µm of dendrite with a range from 1.36 to 3.04 spines/µm in adult human brain. These spine densities in neurons labeled from tissue fixed 10-28 hours postmortem is strikingly consistent with previously reported spine densities of control human CA1 neurons in tissue fixed 2-3 hours postmortem (Merino-Serrais et al., 2013) and to the average CA1 spine density found in non-human primates optimally preserved by perfusion (Leranth, Shanabrough, & Redmond, 2002). To our knowledge, the only other reports of human CA1 spine studies were obtained using two dimensional Golgi methods, which can dramatically underestimate spine density (Liagkouras et al., 2008). Accordingly, studies using Golgi report much lower spine densities (less than 1 spine/µm) (Liagkouras et al., 2008). Therefore, our data from tissue with a 10-28 hours postmortem interval is among the highest reported for human neurons and is comparable to the only existing data from a 2-3 hour postmortem interval. As with all human tissue studies, we cannot rule out that some tissue degradation and spine changes occur with any postmortem interval, but our data strongly suggests that CA1 dendritic spine density is relatively stable over longer postmortem intervals than previously thought.

### 4.3 CA1 spine density and morphology are stable in healthy adult humans

It is well established that cognition and memory naturally declines with normal aging in humans (Buckner, 2004). Unlike in Alzheimer’s and other neurodegenerative diseases, neuron cell loss is not observed in normal, healthy aged brains (Dickstein, Weaver, Luebke, & Hof, 2013). Instead, it is thought that synaptic changes underlie age-related cognitive decline but the nature of the synaptic changes and the brain regions affected remain unclear. The hippocampus is a key brain region that functions during episodic memory but there are surprisingly few studies of hippocampal neuron morphology in humans.

Here, we analyzed spine density, an established proxy for excitatory synapse density (Hering & Sheng, 2001), in hippocampal CA1 neurons in postmortem tissue from healthy subjects ranging in age from 5 months to 71 years old. We found that CA1 apical and basal spine density and morphology are stable throughout adulthood and old age. Though this may seem surprising given the importance of hippocampal CA1 neurons to learning and memory, our results are consistent with studies from rodents and non-human primates. Studies consistently demonstrate that hippocampal dendritic trees, spine density, and excitatory synapse density undergoes little to no change during normal aging in mice, rats, and monkeys (Curcio & Hinds, 1983; Flood & Coleman, 1990; Gonzalez-Ramirez, Velazquez-Zamora, Olvera-Cortes, & Gonzalez-Burgos, 2014; Markham, McKian, Stroup, & Juraska, 2005; Morrison & Baxter, 2012; Pereira et al., 2014; Tsai et al., 2018). Although dendritic spines are relatively stable in CA1, aging may decrease the size of post-synaptic densities in CA1 (Morrison & Baxter, 2012).

In contrast, cortical neurons, and especially those in the prefrontal cortex, have been shown to lose 30 – 60% of their spines and excitatory synapses with normal aging in monkeys (Coskren et al., 2015; Luebke et al., 2015; Morrison & Baxter, 2012; Peters, Sethares, & Luebke, 2008). Thus, it appears that normal aging differentially impacts different brain regions. Neural activity in the prefrontal cortex is necessary for working memory and these cortical neurons are thought to contain more thin spines that are highly flexible and plastic. Therefore, the dynamic nature of prefrontal cortex synapses may make them more susceptible to loss during aging than relatively stable CA1 spines. It is also possible that CA1 synapses undergo age-related changes in spine turnover, presynapse function, or postsynapse function that we cannot detect by analyzing spines at a fixed time point with light microscopy. Electron microscopy has shown that CA1 neurons in aged rats have similar synapse densities than young animals but that those synapses have smaller postsynaptic densities and fewer perforated synapses, a marker of synapse maturity (Geinisman et al., 2004; Nicholson, Yoshida, Berry, Gallagher, & Geinisman, 2004). Moreover, how normal aging impacts inhibitory synapses in the human brain remains unknown. Future studies that analyze synapses in human tissue across the lifespan using electron microscopy could help to resolve these issues. Nonetheless, our study on CA1 spines represents the largest analysis of human CA1 spines to date and because CA1 spines are stable throughout adulthood, our new methodological pipeline should prove useful in identifying changes to CA1 spine density or morphology associated with pathological states including cognitive and mental illnesses or neurodegenerative disorders.

### 4.4 The structure of human mossy fiber boutons changes with age

The DG mossy fiber synapse is a major contributor to pattern separation and spatial memory (Holahan, Rekart, Sandoval, & Routtenberg, 2006; Ramirez-Amaya, Balderas, Sandoval, Escobar, & Bermudez-Rattoni, 2001; Rolls & Kesner, 2016). Mossy fiber synapses connect DG neurons to both CA3 pyramidal neurons and GABA neurons through distinct synaptic elements. Because of their large size, mossy fiber synapses are one of the few types of presynaptic structures that can be easily resolved with light microscopy. Thus, the study of mossy fiber presynapse morphology can provide insights to how the output of the DG may change from a healthy to a diseased state.

In contrast to CA1 spines, we find that mossy fiber presynapse structure significantly changes with age in humans. Specifically, we find that mossy fiber bouton volume, filopodia density, and filopodia length is highest in young children (<2 years) and decreases throughout adulthood, while bouton volume and filopodia density may increase again in very old age. High numbers of mossy fiber filopodia are also observed in young rodents (Martin et al., 2015; Wilke et al., 2013) and may reflect a developmental phenomenon with numerous motile filopodia still undergoing synapse formation (Tashiro, Dunaevsky, Blazeski, Mason, & Yuste, 2003). In young mice, up to 50% of the mossy fiber filopodia do not have a synapse (Martin et al., 2015) and these synapse-lacking filopodia may be preferentially lost to reach adult numbers.

Mossy fiber filopodia are known to be dynamic structures and temporarily increase in density after learning in adult mice (Ruediger et al., 2011). Therefore, mossy fiber filopodia may be influenced by neural activity or homeostatic mechanisms. Moreover, age-related spatial memory decline has been postulated to be mediated by increased excitability of CA3 neurons due to decreased GABAergic signaling or loss of GABAergic interneurons (Thome, Gray, Erickson, Lipa, & Barnes, 2016; Yassa et al., 2011). Because mossy fiber filopodia synapse primarily with GABAergic interneurons, it is possible our observed increase in mossy fiber filopodia density with old age reflects an attempt by a normal healthy brain to rebalance network activity. Conversely, increased mossy fiber filopodia in old age could represent a pathological response of aging that contributes to memory decline. Consistent with this hypothesis, a mouse model of familial Alzheimer’s disease also has increased density of mossy fiber filopodia compared to control mice (Wilke et al., 2014). It will be interesting to follow up this study with analyses of mossy fiber structures in brain tissue from Alzheimer’s disease patients. Taken together, our study presents the first structural analysis of mossy fiber synapses in the human brain and suggests that mossy fiber synapses, in contrast to CA1 synapses, are particularly dynamic during aging.

In conclusion, our proposed DiI-labeling is applicable for accurate and higher throughput analysis of dendritic spines and mossy fiber boutons in human tissue samples with broad postmortem intervals. Using this method, it will be possible to study the alterations in pre- and postsynaptic structures throughout the brain in disease-specific human postmortem tissue samples.

## Acknowledgments and Funding

This work was supported by grants from the National Institute of Mental Health R01MH099134 (HC) and R01MH105426 (MEW) and the University of Utah Medical School Psychiatry Department Research Fund (SCD). We especially thank the Utah State Office of the Medical Examiner staff who made this study possible and members of the Williams and Coon labs for reading the manuscript.

